# Dissecting the genetic basis of a complex *cis*-regulatory adaptation

**DOI:** 10.1101/029207

**Authors:** Santiago Naranjo, Justin D. Smith, Carlo G. Artieri, Mian Zhang, Yiqi Zhou, Michael E. Palmer, Hunter B. Fraser

## Abstract

Although single genes underlying several evolutionary adaptations have been identified, the genetic basis of complex, polygenic adaptations has been far more challenging to pinpoint. Here we report that the budding yeast *Saccharomyces paradoxus* has recently evolved resistance to citrinin, a naturally occurring mycotoxin. Applying a genome-wide test for selection on *cis*-regulation, we identified five genes involved in the citrinin response that are constitutively up-regulated in *S. paradoxus.* Four of these genes are necessary for resistance, and are also sufficient to increase the resistance of a sensitive strain when over-expressed. Moreover, *cis*-regulatory divergence in the promoters of these genes contributes to resistance, while exacting a cost in the absence of citrinin. Our results demonstrate how the subtle effects of individual regulatory elements can be combined, via natural selection, into a complex adaptation. Our approach can be applied to dissect the genetic basis of polygenic adaptations in a wide range of species.

**Author Summary:** Adaptation via natural selection has been a subject of great interest for well over a century, yet we still have little understanding of its molecular basis. What are the genetic changes that are actually being selected? While single genes underlying several adaptations have been identified, the genetic basis of complex, polygenic adaptations has been far more challenging to pinpoint. The complex trait that we study here is the resistance of *Saccharomyces* yeast to a mycotoxin called citrinin, which is produced by many other species of fungi, and is a common food contaminant for both humans and livestock. We found that sequence changes in the promoters of at least three genes have contributed to citrinin resistance, by up-regulating their transcription even in the absence of citrinin. Higher expression of these genes confers a fitness advantage in the presence of citrinin, while exacting a cost in its absence— a fitness tradeoff. Our results provide a detailed view of a complex adaptation, and our approach can be applied to polygenic adaptations in a wide range of species.

## Introduction

Historically, most studies pinpointing the genetic basis of polymorphic traits have focused on protein sequence changes of large effect, because these have been the most amenable to identification. For example, thousands of coding region mutations have been implicated in human diseases with Mendelian inheritance [1]. In contrast, non-coding mutations and polygenic traits have traditionally received far less attention.

However this situation has radically changed with the advent of genome-wide association studies (GWAS). In these studies, millions of SNPs can be tested for statistical association to any trait of interest. Two clear patterns have emerged from hundreds of human GWAS: most traits are highly polygenic, and most associations are in non-coding regions that are not in linkage disequilibrium with protein-coding changes (and thus cannot be acting via changes in protein sequence) [2-4]. For example, out of 697 loci associated with height, 86% are non-coding [5]. This suggests that natural selection will overwhelmingly result in polygenic *cis*-regulatory adaptations, composed of variants with very small individual effects, since selection acts on whatever heritable variation is available. Indeed, thousands of loci are involved in selection on height in Europeans, a clear example of polygenic adaptation [6]. Moreover, recent genome-wide comparisons of the proportions of adaptations in coding vs. non-coding regions in humans and sticklebacks support the prevalence of non-coding adaptations [7-9].

Two methods have successfully identified the genes underlying regulatory adaptations. One is QTL/association mapping, which has led to several beautiful examples of single-locus adaptations [10]. However QTL mapping in most species is only practical for loci of large effect, and thus is not well-suited for studying the evolution of complex traits, which are by definition polygenic.

The second method is the “sign test” framework that we and others have developed [11-12]. The goal of this approach is to identify cases where selection has led to up- or down-regulation of multiple genes via independent mutations. First the *cis*-regulatory divergence between two species is quantified genome-wide via allele-specific expression (ASE) analysis in an F1 hybrid [11]. This results in directionality information for every gene (e.g. for gene X, the species A allele is up-regulated compared to the species B allele). Any group of genes whose expression is evolving under the same selection pressure in the A and B lineages should have a similar frequency of A alleles up-regulated as in the entire genome. For example if 50% of genes with ASE have A allele up-regulation, then any random subset of these should have roughly 50% as well. If a strong deviation from 50% is detected for some gene set—e.g. all 20 genes in a pathway have the A alleles up-regulated—then this indicates the action of lineage-specific selection. Importantly, this approach does not make many assumptions of other methods for detecting selection such as constant population size, lack of epistasis, or neutrality of synonymous sites [11]. And unlike approaches that can rank genes but cannot determine which (if any) of them are inconsistent with neutrality (such as F_ST_, iHS, and the ratio of expression divergence between species to expression diversity within species), the sign test's null model allows for confident identification of gene sets under lineage-specific selection [11]. Finally, the sign test is most powerful when many genes are involved, making it uniquely well-suited for studying complex traits.

In the current work, we applied the sign test to ASE data from a hybrid between two species of budding yeast, *Saccharomyces cerevisiae* and *S. paradoxus* (specifically the reference strains S288c and CBS432, hereafter abbreviated *Sc* and *Sp*; note that these abbreviations refer specifically to the two reference strains, and not the two species as a whole). The results led us to focus on a specific mycotoxin (a toxin produced by fungi) called citrinin. Citrinin is produced by a number of Ascomycota fungi, including several species in the *Aspergillus, Penicillium,* and *Monascus* genera. It increases mitochondrial membrane permeability and causes oxidative damage via an unknown mechanism, and is a potent nephrotoxin in mammals [13-14]. Because citrinin is toxic to yeast growing on fermentable carbon sources, which do not require mitochondria, toxicity is likely caused by oxidative damage to other cellular components [13]. The transcriptional response may be an important means to mitigate citrinin toxicity, and in *Sc,* hundreds of genes are induced or repressed in response to even a low level of citrinin [15]. Although citrinin is a common food contaminant, making it a major health concern for both humans and livestock [13-14], and has been the subject of hundreds of publications, the evolution of citrinin resistance has not been previously investigated.

## Results

### A sign test reveals select/on on the cis-regulation of citrinin-induced genes

In an effort to identify groups of genes subject to lineage-specific selection, we applied a sign test (described above) to allele-specific gene expression levels from an *Sc/Sp* hybrid grown in YPD (rich glucose) media [16]. Our objective was to identify any gene sets (such as pathways or other functionally related groups) with a biased directionality of ASE, implying the action of lineage-specific natural selection [11]. Testing a collection of publicly available gene sets (see Methods), we found only one set with a highly significant bias: genes induced by exposure to citrinin, a naturally occurring mycotoxin, were over-represented among genes with *Sp*-biased ASE. Of 11 genes that were reported to be induced at least 10-fold in response to citrinin (in *Sc*) [15], four were among the top 1% of *Sp*-biased ASE genes (Fig. 1; hypergeometric *p* = 2.7 × 10^−6^). In fact, these four genes included the first and third most *Sp*-biased genes in the entire genome. In other words, *Sp* alleles have markedly higher expression of several genes that are induced in *Sc*’s citrinin response, even though ASE was measured in the absence of citrinin [16].

**Figure 1.**
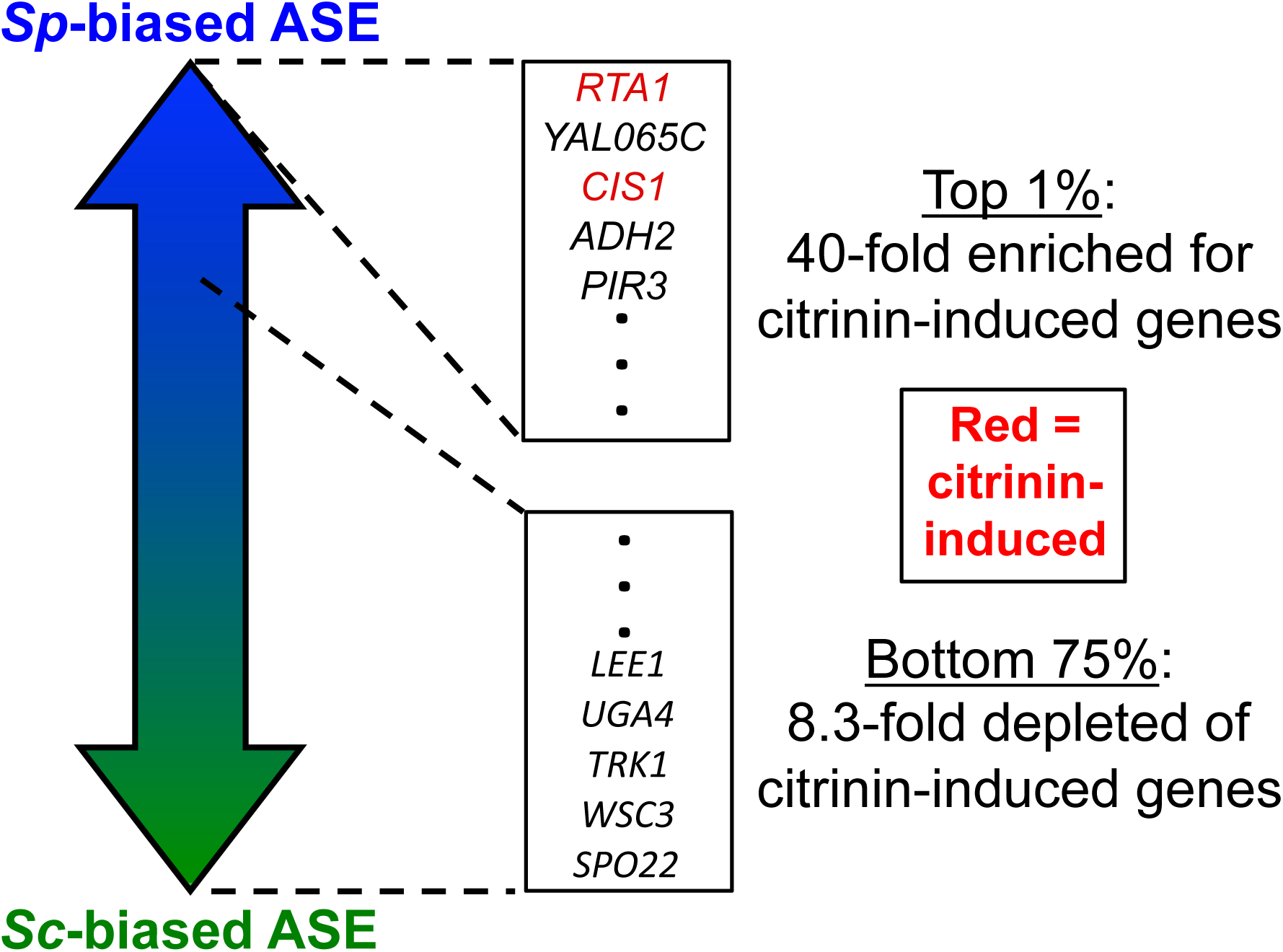
Results of the sign test for selection on *cis***-**regulation. For a description of the test and its results, see the main text and Methods. Note that only two citrinin-induced genes are shown, but four were present in the top 1% of Sp-biased genes, and ten in the top 25%.

In contrast, none of the 11 citrinin-induced genes had even slightly *Sc*-biased ASE in the *Sc/Sp* hybrid. All 11 had some degree of *Sp*-biased ASE, and 10/11 were in the top 25% of *Sp*-biased genes (Fig. 1; *p* = 7.8 × 10^−6^). The complete absence of citrinin-induced genes among *Sc*-biased genes, taken together with their 40-fold enrichment among the most strongly *Sp*-biased genes, is not consistent with a neutral model of gene expression evolution leading to random directionality of ASE [11]; instead, it suggests that lineage-specific selection shaped the *cis*-regulation of these genes.

### Sp *has recently evolved resistance to citrinin*

The results of our sign test (Fig. 1) suggested that *Sp*’s constitutive up-regulation of *Sc’*s citrinin-induced genes may confer a greater resistance to citrinin. To test this, we measured the growth of *Sc* and *Sp* in three concentrations of citrinin: 0 ppm (parts per million), 300 ppm, and 600 ppm. At 300 ppm, *Sc* showed a clear shift towards slower growth, while *Sp* was essentially unperturbed (Fig. 2A-B). At 600 ppm, *Sp* was affected, but not nearly as strongly as *Sc;* the effect on *Sp* at 600 ppm was roughly similar to that on *Sc* at 300 ppm. These results are consistent with our hypothesis that *Sp* would show increased resistance.

**Figure 2.**
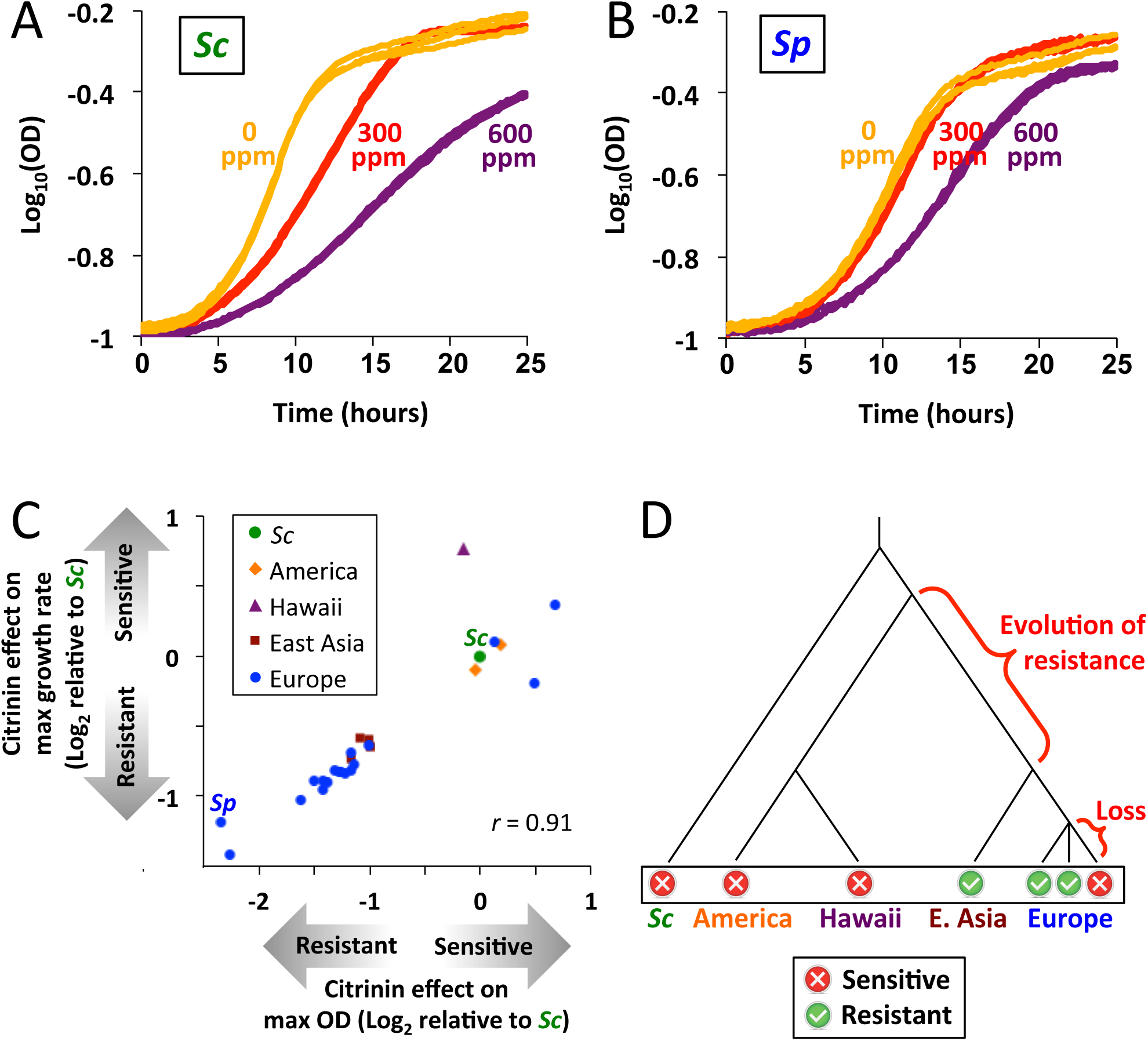
*Sp* recently evolved resistance to citrinin. **A.** *Sc* growth at different concentrations of citrinin. Three replicates are shown for each condition. **B.** *Sp* growth at different concentrations of citrinin. Three replicates are shown for each condition. **C.** Two metrics of resistance, maximum growth rate and maximum cell density, are shown for 25 *S. paradoxus* strains and *Sc.* Results are plotted as log-ratios relative to *Sc.* **D.** Citrinin resistance treated as a binary trait and plotted on a phylogenetic tree [44]. The most parsimonious evolutionary scenario is shown in red text. Branch lengths are not to scale.

To investigate the evolutionary history of the resistance phenotype, we measured growth rates and cell densities at saturation (“max OD”) for a panel of 25 diverse *S. paradoxus* strains. These strains belong to four major clades, defined on the basis of partial genome sequences [17], which also reflect their geographic origins: European (including Western Russia), East Asian (Eastern Russia and Japan), Hawaiian, and American. We also included *Sc* as an outgroup. Across all strains, the effect of citrinin on growth rate and on max OD was highly correlated (Pearson *r* = 0.91; Spearman *r* = 0.96), and strains fell into distinct clusters based on their sensitivities (Fig. 2C). The most sensitive group included *Sc,* the Hawaiian strain, both American strains, and 3/18 European strains. In contrast, the resistant group consisted of all four East Asian strains, and 15/18 European strains (including the two most resistant strains, *Sp* and Q62.5).

Representing resistance as a binary trait on the *S. paradoxus* phylogeny, the most parsimonious explanation is that resistance evolved after the split between the Hawaiian/American clade and the European/East Asian clade, and was then lost in a subset of the European strains (Fig. 2D) (this loss could have occurred via new mutations, or admixture from a sensitive *S. paradoxus* strain; see Discussion). Any other scenario would require multiple independent gains and/or losses. These results indicate the difference between *Sc* and *Sp* (Fig. 2A-B) is likely due to resistance being gained in a recent ancestor of *Sp,* as opposed to being lost in the *Sc* lineage.

To determine whether citrinin resistance represents a more general pleiotropic trait—such as resistance to many different toxins—we compared our growth data (Fig. 2C) to growth rates of the same strains in 200 diverse conditions [18]. These include several oxidative stress agents (aminotriazole, paraquat, dithiothreitol, CdCl_2_, and CoCl_2_) and dozens of other toxins. None of these showed a similar pattern of resistance across strains as we observed for citrinin: the maximum correlation across all 200 conditions was *r* = 0.44 (*n* = 23 strains; not significant after correction for 200 tests). In fact, *Sp* showed the *least* resistance to all five oxidative stress conditions (among 22 *S. paradoxus* strains and *Sc*), the opposite of our observation for citrinin. Together, these results suggest that citrinin resistance does not represent a more general resistance to toxins or oxidative stress.

### Candidate genes revealed by RNA-seq in hybrids

To further characterize the effect of citrinin on gene expression, we performed RNA-seq on *Sc/Sp* hybrid yeast exposed to 600 ppm citrinin, as well as in rich media lacking citrinin for comparison. This allowed us to measure the effect of citrinin on each gene’s ASE, and thus to measure the *cis*-regulatory contributions of each species to the induction or repression of each gene. We found strong agreement between our biological replicates (*r* = 0.96-0.99; S1 Fig), and moderate concordance with published microarray data from the *Sc/Sp* hybrid [16] and *Sc’*s response to citrinin [15] (S2 Fig).

To determine if the transcriptional response to citrinin was species-specific, we analyzed the responses of *Sc* and *Sp* alleles separately in our hybrid RNA-seq data. We observed highly concordant responses to citrinin (Fig. 3A). In fact, among 114 genes with at least 3-fold induction or repression for alleles from both species, all of them responded to citrinin in the same direction for both (*r* = 0.96 for these genes). As a result, ASE is broadly similar in rich media and in citrinin, as has been reported for the *Sc/Sp* hybrid in other conditions [16].

**Figure 3.**
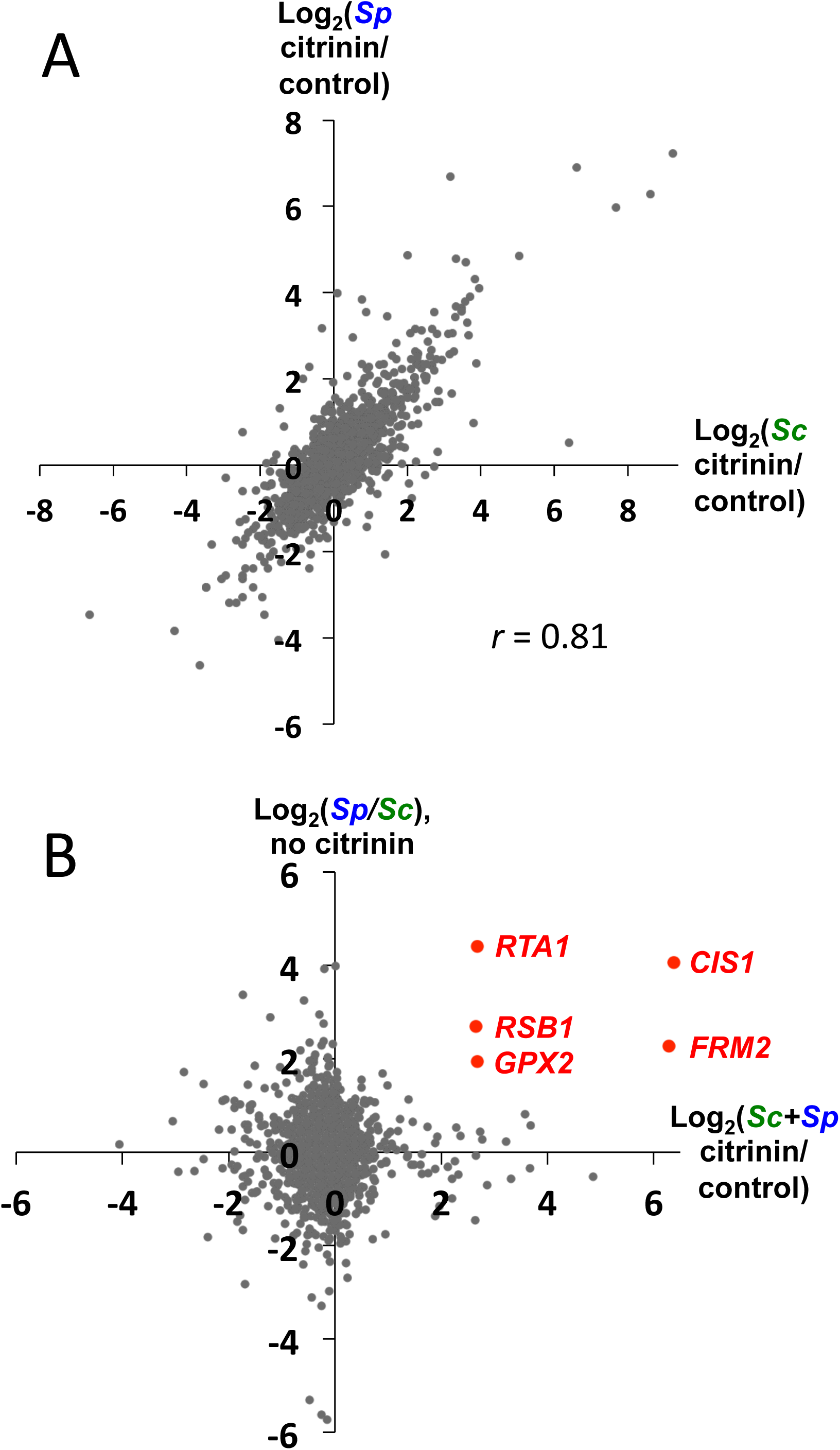
RNA-seq reveals candidate genes. **A.** Citrinin has largely similar effects on *Sc* alleles and *Sp* alleles in the *Sc/Sp* hybrid; i.e. ASE is similar with or without citrinin. **B.** Candidate genes were identified based on having strong *Sp*-biased ASE (in YPD; FDR < 0.05) [16], and strong up-regulation in response to citrinin (of both *Sc* and *Sp* alleles; the smaller of these two in our RNA-seq data is plotted; binomial *p* < 10^−5^ for each biological replicate of each allele).

To identify candidate genes that may be contributing to the polygenic selection that we detected (Fig. 1), we selected those with the strongest combination of citrinin-induction and *Sp*-biased ASE. To visualize this, we plotted each gene’s citrinin response against *Sp/Sc* ASE in the absence of citrinin (Fig. 3B). Requiring at least 4-fold citrinin induction and 4-fold *Sp*-biased ASE, we identified four genes; a fifth gene that was slightly below the ASE cutoff was included as well (Fig. 3B, red points). These constituted our top candidates for genes that may be involved in the evolution of citrinin resistance in *Sp.* Notably, no genes had both a 4-fold ASE bias and citrinin-response in any of the other three possible pairs of directions (the three additional quadrants in Fig. 3B), indicating the rarity of this combination.

If citrinin resistance evolved after the split of European/East Asian and American/Hawaiian strains (Fig. 2D), we may expect that genes involved in this adaptation would show an ASE bias towards *Sp* (European) alleles in hybrids between these two clades, as we observed for *Sc/Sp.* To test this, we performed RNA-seq in a hybrid between *Sp* and an American *S. paradoxus* strain (DBVPG6304). We could assess ASE for 4/5 candidate genes; three of these showed Sp-biased ASE (*RTA1,* binomial *p* = 2×10^−4^; *FRM2, p* = 0.05; *CIS1, p* = 8×10^−8^), while the fourth had strong American-biased ASE (*RSB1, p* = 3×10^−55^). Therefore our prediction of ASE directionality was validated for 3/4 candidate genes.

### Characterization of the candidate genes

As an initial test of these five candidate genes, we deleted each gene individually from *Sp* and tested the effect on citrinin resistance (all five were nonessential in YPD). For four of the five gene deletions, *Sp* resistance was significantly decreased (Fig. 4A); the only exception was *RSB1,* the same gene that did not show *Sp*-biased ASE in the *Sp*/American hybrid. These four genes are involved in a variety of functions: a mitochondrial glutathione peroxidase involved in the oxidative stress response (*GPX2*), an oxidoreductase also involved in oxidative stress response (*FRM2*), a lipid-translocating exporter of the plasma membrane (*RTA1*), and a mitochondrial protein of unknown function (*YLR346C,* which we rename as *CIS1,* for “<u>CI</u>trinin <u>S</u>ensitive knockout”). The functions of all four genes are consistent with a role in citrinin resistance: three are involved in mitochondria/oxidative stress, processes directly related to citrinin [13-14]; and the fourth, *RTA1,* has been implicated in toxin resistance (*Sc* strains missing *RTA1* are highly sensitive to a mycotoxin called myriocin [19], and over-expression confers resistance to multiple toxins [20]). Considering the four lines of evidence converging on these genes—induction in response to citrinin, constitutive up-regulation of their *Sp* alleles (in two different hybrids), the effects of their knockouts on *Sp* citrinin resistance, and functional annotations—we focused further efforts on these four candidates.

**Figure 4.**
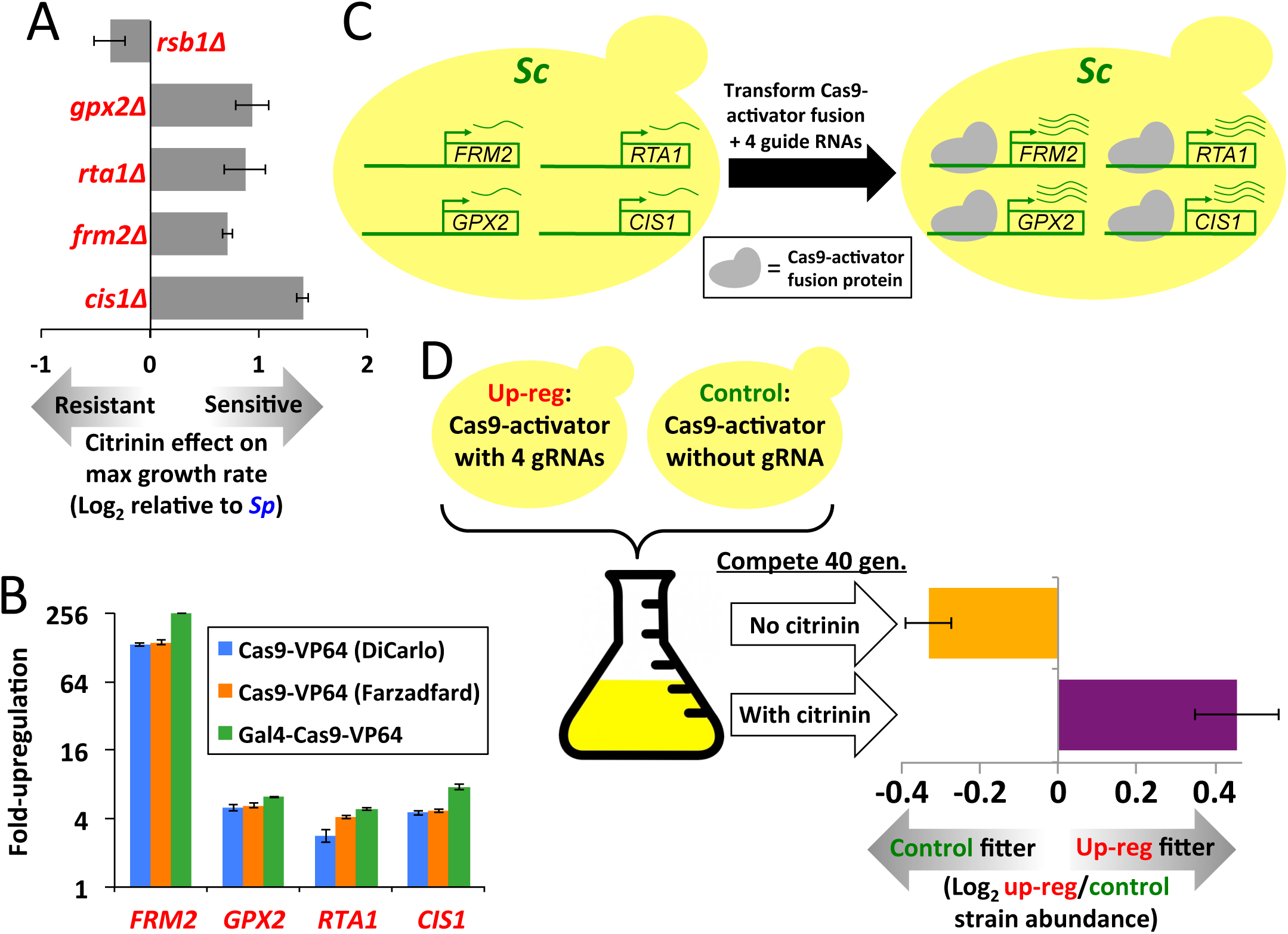
Effects of candidate gene deletion and over-expression. **A.** The five candidate genes were deleted from *Sp,* and the effect on resistance to 300 ppm citrinin was measured. Four deletions increased sensitivity (t-test *p* < 0.005 for each), and were studied further. Error bars show 1 S.E. **B.** The effects of three different dCas9 fusion proteins targeted to four candidate genes were measured by qPCR. **C.** Illustration of our strategy to over-express four genes via dCas9 fusion protein. **D.** We used direct competition for 40 generations to measure the relative fitness of the four-gene over-expression strain (with a Gal4-dCas9-VP64 dual fusion) vs. a control strain containing the same plasmid, but lacking any gRNAs. Conditions were YPD, and YPD + 300 ppm citrinin. Error bars show 1 S.E.

A complement to testing whether gene deletion leads to trait loss is to test whether over-expression leads to trait gain. To explore this, we used the CRISPR/Cas9 system, in which the Cas9 protein can be directed to a specific genomic site by use of a guide RNA (gRNA) complementary to the target DNA site. When a transcriptional activation domain is fused to a nuclease-dead mutant of Cas9 (called dCas9), the resulting protein acts as a strong activator of its target genes [21]. This is an attractive system for simultaneous over-expression of multiple genes, since multiple guide RNAs can be delivered to cells on a single plasmid.

To test the efficacy of this system, we compared the induction levels of our four candidate genes using three different dCas9-activator fusion genes. One was a previously published fusion to VP64 (a domain derived from a strong viral transcription factor) [21]; the second was a dCas9-VP64 fusion that we created using a dCas9 gene with an alternative codon optimization; and the third was a dual-fusion that we created with VP64 at the C terminus and the Gal4 activation domain at the N terminus. We found that all three dCas9 constructs could induce all four genes, though the level of induction varied substantially between genes, with *FRM2* achieving far higher induction than the other three (Fig. 4B), perhaps due to its lower basal expression level. Comparing the three dCas9 fusion proteins, we observed similar induction levels for all three, though our dual-fusion construct achieved slightly (1.2-1.8-fold) higher induction for all four genes. We then used this dCas9 dual-fusion gene to over-express all four genes in a single strain (Fig 4C).

To measure the effect of over-expression on fitness, we performed direct competition between the over-expression strain vs. a control (a strain with a dCas9 plasmid lacking any gRNAs), for 40 generations (Fig 4D). We incorporated 6-base barcodes into each plasmid, and sequenced these before and after each competition to estimate strain abundances; each strain was tagged by three different barcodes, to reduce any barcode-specific biases. In the absence of citrinin, the over-expression strain had a small but consistent (across 12 replicates) fitness disadvantage of 0.6% per generation (resulting in 20% lower abundance than the control after 40 generations). In contrast, in 300 ppm citrinin, the over-expression strain had a 0.8% fitness advantage (37% higher after 40 generations; t-test p = 3×10^−5^ comparing conditions). These results suggest that over-expression of these genes leads to a condition-specific fitness tradeoff.

### Individual promoters contribute to citrinin resistance

*Cis*-regulatory divergence in mRNA levels can be caused by changes at either the transcriptional or post-transcriptional (e.g. mRNA stability) level. We hypothesized that the major causal mutations may have affected transcription via changes in promoter sequences between *Sc* and *Sp.* To test this, we replaced the promoter region (~1 kb of noncoding DNA upstream of each gene) of each of our four candidate genes in *Sc* with the orthologous promoter from *Sp* (Fig. 5A), using an approach that leaves no foreign DNA behind [22]. Boundaries were chosen to overlap stretches of perfect *Sc/Sp* conservation to ensure correct placement of the *Sp* sequence in the *Sc* genome, as well as to facilitate homologous recombination at the target site.

**Figure 5.**
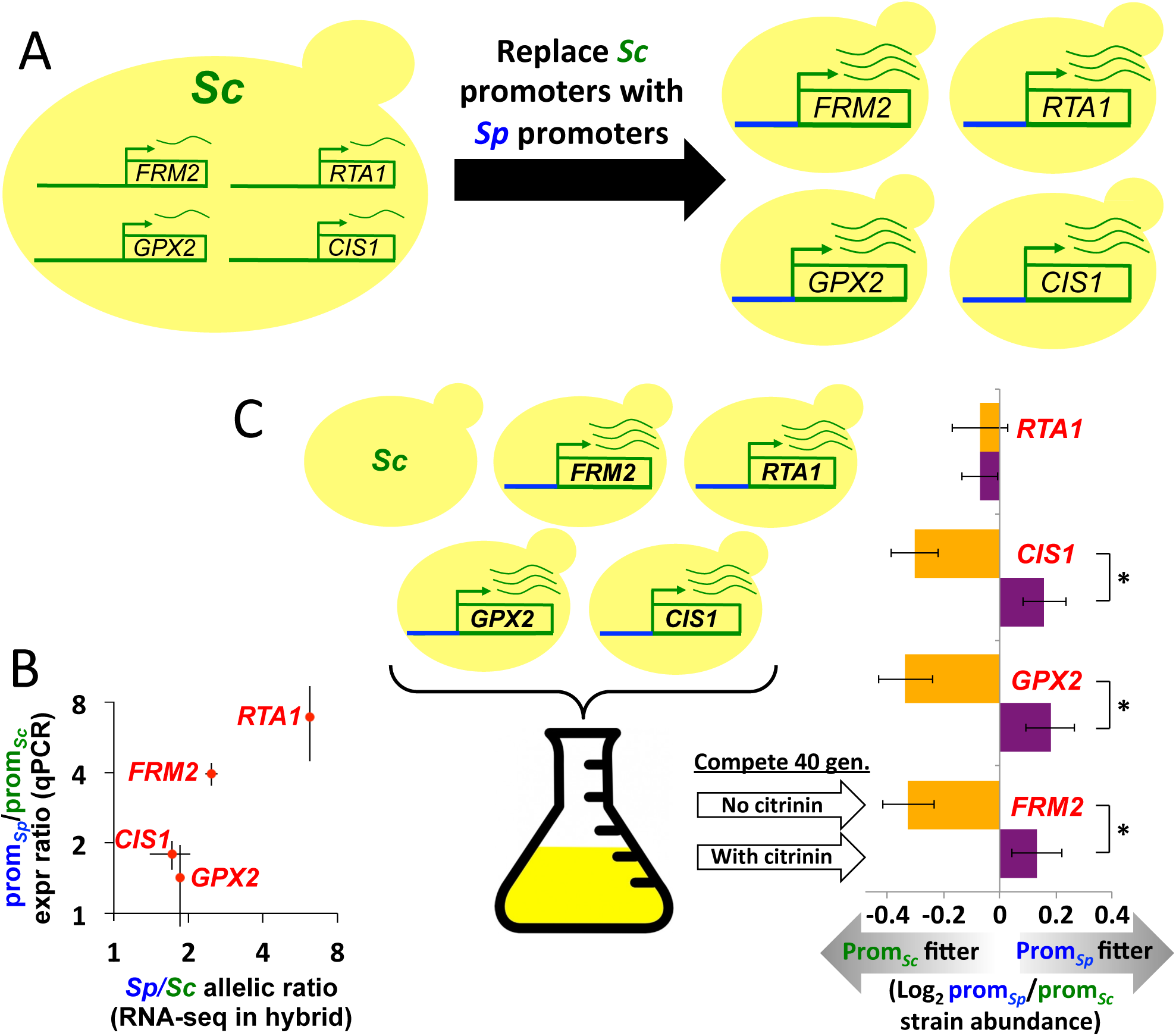
Promoter replacement reveals contributions of individual regulatory regions. **A.** Illustration of our promoter replacement strategy. Green promoters on the left are from *Sc,* blue promoters on the right are from *Sp.* **B.** RNA-seq in the *Sc/Sp* hybrid, which measures the overall *cis*-regulatory divergence between these strains, is in approximate agreement with the effect of the promoter replacements, measured by qPCR (*r* = 0.92). Error bars show 1 S.E. **C.** After competitive growth for 40 generations, promoter replacement strains for 3/4 candidate genes show a similar pattern of fitness advantage in the presence of 300 ppm citrinin, and disadvantage in its absence. Error bars show 1 S.E.

In each “promoter-replacement” strain we performed quantitative PCR on the downstream gene, to assess the effect on mRNA levels. We found between 1.4 and 6.9-fold up-regulation of the four *Sp* promoters, compared to the orthologous *Sc* promoters. Comparing these results to the total *cis*-regulatory divergence based on our RNA-seq data in the *Sc/Sp* hybrid (Fig. 2A), we found good concordance (Fig. 5B, *r* = 0.92). This suggests that the promoters are likely responsible for most, if not all, of the *cis*-regulatory divergence for these genes between *Sc* and *Sp.*

To test the fitness effects of this *cis*-regulatory divergence, we pooled all four promoter replacement strains together with their *Sc* parent, and grew them in direct competition. We observed a similar pattern for 3/4 candidate genes, with the *Sp* promoter leading to 0.2-0.3% higher fitness in the presence of citrinin, but ~0.6% lower fitness in its absence (Fig 5C; 18 replicates per condition; t-test *p* < 0.01 for each). The fourth gene, *RTA1,* showed no detectable fitness difference in the two growth conditions (*p* = 0.51). These results suggest an overall similar condition-specific fitness tradeoff of the natural *cis*-regulatory divergence as we observed for the 4-gene over-expression strain, with each of three promoters contributing a small amount to citrinin resistance.

## Discussion

In this work, we have discovered a polygenic *cis*-regulatory adaptation, and investigated its genetic basis and effect on fitness. Our results establish that changes in the promoters of at least three genes have contributed to *Sp’*s recently evolved resistance to citrinin, which has a fitness cost in the absence of citrinin. The role of these genes in *Sp’*s citrinin resistance is supported by six lines of evidence: induction in response to citrinin, constitutive up-regulation of their *Sp* alleles (compared to *Sc* and an American strain of *S. paradoxus*), functional annotations, decreased resistance via gene deletion in *Sp*, increased resistance via over-expression in *Sc,* and the fitness effects of promoter-replacements.

Isolating the effects of individual promoter regions has allowed us to gain a deeper understanding of how distinct loci can contribute to polygenic adaptation. For example, the fitness cost of increased expression in the absence of citrinin may explain why these genes are induced in response to citrinin, as opposed to being expressed at constitutively high levels. Despite this cost, *Sp* does express all four genes at somewhat higher levels than *Sc* even without citrinin present (Fig 3B), which may confer a net benefit if citrinin is encountered more often by *Sp* than by *Sc.* Alternatively, the fitness costs we observed in *Sc* may be reduced in *Sp* by compensatory changes at other loci.

Having identified this gene expression adaptation, a natural question is what selective pressure(s) caused it. Although citrinin represents a plausible candidate, since it is a widespread naturally occurring toxin, it is also possible that something else (e.g. another toxin) was the actual selective agent, with citrinin resistance being a pleiotropic side-effect. Unfortunately there is no experiment that could unambiguously identify the selective agent, since pleiotropic effects of unknown/unmeasured traits are always a possibility; however our finding that citrinin resistance does not correlate with fitness across 200 other growth conditions [18] suggests that it is not a highly pleiotropic trait.

An interesting aspect of our data is the strong correlation between maximum growth rate and maximum cell density across wild isolates (Fig. 2C). Similar correlations (*r* = 0.85 and 0.77) were also observed between these same two variables among other strains of yeast [18,23]. Ibstedt et al. [23] proposed that the correlations between fitness components of natural isolates are not due to pleiotropic effects, but rather that selection has fixed multiple variants affecting different fitness components in particular lineages. In other words, the presence of a correlation between fitness components may, by itself, be evidence of selection on condition-specific fitness [23]. In the case of citrinin resistance in *Sp,* this interpretation is quite consistent with our detection of polygenic selection via the sign test.

Another intriguing property of the resistance across *S. paradoxus* isolates is that resistance is roughly bimodal (Fig. 2C). This makes sense if resistance evolved once, and has not changed drastically since then in most of the strains tested here. The exception to this is the three European strains that have lost their resistance, perhaps due to new mutations and/or variants introduced via admixture with sensitive strains. Whether resistance was lost once or multiple times among European strains would be difficult to determine—although these three strains do not cluster together within the European phylogeny [17], this represents only a genome-wide average phylogeny that many loci, including those involved in the loss of citrinin resistance, may not follow.

Our inference that resistance evolved within *S. paradoxus* (Fig. 2D) is based on parsimony, i.e. explaining a phylogenetic pattern with the fewest possible transitions. However if loss of citrinin resistance is more likely than gain then a scenario with an ancestral state of resistance, followed by three loss events (one in *Sc* and two in *S. paradoxus*), could be more plausible. If this were the case, the lineage-specific selection that we detected with the sign test would more likely represent selection for lower expression of these genes in *Sc,* to avoid the fitness cost of their expression in the absence of citrinin (Fig. 4D and 5C). A relaxation of selection (i.e. loss of constraint) is another possible explanation for down-regulation [11], though in this case would not be consistent with either the fitness effects of promoter-replacements (which are large enough to be strongly selected), or the correlation of fitness components (Fig. 2C) that suggests selection is acting on citrinin resistance within *S. paradoxus* [23]. Therefore regardless of the ancestral state, the differences we have observed between *Sc* and *Sp* are likely to be adaptive.

In this work, we have only explored one facet of this polygenic adaptation, namely *cis*-regulatory divergence of mRNA levels in a specific environment (YPD media at 30°C). Many other types of evolutionary changes may also be involved, such as changes in protein sequences, *trans*-acting factors, translation, etc. Moreover, it is quite likely that the *cis*-regulatory divergence of additional genes also contributes to *Sp*’s citrinin resistance; for example in our initial analysis (Fig. 1), there were five genes with strong citrinin-induced upregulation and moderate *Sp*-biased ASE (in the top 25% of genes) that we did not pursue, as well as many more genes with weaker supporting evidence. Consistent with the idea that additional loci likely contribute, the four promoter replacements we tested cannot account for the large difference in resistance between *Sc* and *Sp* (Fig 2A-B) if we assume additivity of the promoter-replacement fitness effects and equivalence of the fitness measurements in these two experiments.

For regulatory changes of *RTA1,* for which we could not detect an effect on citrinin resistance, many possibilities exist—e.g. condition-specific effects (gene-by-environment interaction), or genetic background-specific effects (gene-by-gene interaction, i.e. epistasis), or effects that were too small to measure, or it may have no effect at all. It is clear that even for this one adaptation (as well as every other polygenic adaptation), we are still far from having a complete understanding.

Looking ahead, an important question is what approaches will be most useful for understanding the genetic basis of evolutionary adaptations. QTL mapping has been very effective in localizing large-effect loci down to specific genomic regions, and has led to the identification of several genes underlying adaptations [10]. However it does have limitations. First, an adaptive phenotype must be identified before starting QTL mapping; though in many cases, such as citrinin resistance, we do not know ahead of time which traits may be adaptive. Second, QTL mapping is generally only practical for loci of large effect (with the exception of pooling-based approaches in yeast [24]), which may be rare among adaptations, given the highly polygenic nature of most traits [2-4]. Third, specific genes are not implicated; QTL peaks typically contain dozens or even hundreds of genes, of which only one may be involved in the trait. And fourth, even for loci of large effect, QTL mapping requires generating, genotyping, and phenotyping hundreds of F2 individuals to result in a reasonably sized QTL peak.

In contrast, the sign test approach we have employed allows us to circumvent these limitations. First, no adaptive phenotype must be known a priori. Second, loci of very small effect can be identified. Third, specific genes are implicated. And fourth, no F2 crosses are required; a single F1 is sufficient to measure all *cis*-regulatory divergence via RNA-seq. Of course, the sign test is limited to polygenic *cis*-regulatory adaptations in which one direction of change (up- or down-regulation) dominates; but for these, which may be quite frequent (see Introduction), the sign test can rapidly identify high-confidence candidate genes. Finally, the sign test can be applied to a wide range of species, including fungi, plants, and metazoans [25-31].

A major limitation of the sign test has been that it has previously relied on functional annotations of genes, thus limiting it to species with reasonably well-annotated genomes. However in this work, we found that gene sets and specific candidate genes can be defined on the basis of gene expression data alone (such as the citrinin-response data of ref. 15)—an important advance for future applications of this approach to non-model organisms. Combined with other recent advances in genetic tools, such as CRISPR/Cas9, we believe that pinpointing specific genes and genetic variants underlying complex, polygenic adaptations is now readily achievable across a wide range of species.

## Methods

### Sign test for selection on cis-regulation

Allele-specific expression data from the *Sc/Sp* hybrid in YPD were obtained from [16]. Two genes with strong *Sp*-biased ASE due to being used as auxotrophic markers in *Sc* (*URA3* and *MET17*) were removed, leaving 4394 measured genes. The top 1%, 5%, and 25% of genes with the strongest ASE in each direction were submitted to FunSpec, a tool for gene set enrichment analysis [32]. The top enrichment at all three cutoffs for *Sp*-biased ASE (Hypergeometric *p* < 0.01 after Bonferroni correction for multiple tests) was the Gene Ontology set “response to toxic substance.” Examination of the enriched genes revealed that their annotations were all derived from a single publication, measuring the transcriptional response to citrinin [15]. Thus subsequent analysis (Fig. 1) was performed using these induced genes as the set of interest. Induced genes were defined as those with at least 10-fold average induction (Table 1 of ref. 15), excluding four paralogous genes found to have cross-hybridization [15]. Out of 17 induced genes, 11 had ASE measurements [16] (In decreasing order of *Sp/Sc* ASE ratio from ref. 16: *RTA1, CIS1, RSB1, FRM2, ECM4, FLR1, AAD4, GRE2, GTT2, YML131W,* and *YLL056C*). None of these 11 genes were next to one another in the genome, so their ASE was likely caused by independent *cis*-regulatory changes, as required by the sign test [11]. No significant ASE directionality bias was observed for citrinin-repressed genes.

### Fitness assays

To perform quantitative growth rate measurements (Fig 2A-C and 4A), we grew strains in 96-well plates and measured OD600 at 12-minute intervals using an automated plate reader (Tecan) until cultures reached saturation. Experiments were performed at 30°C in YPD media [33] (with or without citrinin). Citrinin (Santa Cruz Biotechnology) was dissolved in DMSO at a concentration of 20,000 ppm, prior to addition to growth media. An equal amount of DMSO was added to the no-citrinin controls, as was in each matched experiment with citrinin+DMSO. All growth assays were performed with each strain distributed across the plate (e.g. alternating rows/columns for the different strains/conditions) to minimize any spatial variation in conditions across the plate.

Maximum growth rate was estimated by performing a linear regression on log_10_(OD) values vs. time, for every set of 20 consecutive time points (4 hours), and the highest slope was recorded. Maximum OD was calculated across all time points. For clarity of presentation, we show growth rates as log-ratios relative to a fixed reference (*Sc* in Fig. 2C, and *Sp* in Fig. 4A). For example in Fig. 4A, a strain with a 2-fold greater reduction in max growth rate than *Sp* after citrinin exposure would have a value of 1; a 4-fold lower reduction in max growth rate than *Sp* would have a value of -2.

### RNA-seq and ASE analysis

An *Sc/Sp* diploid hybrid yeast strain was produced by mating *Sc* (BY4716: MATα lys2 ura3::KAN) and *Sp* (CBS432: MATa ura3::HYG), followed by selection on plates containing hygromycin B and G418 (Sigma). Four single colony picks of the hybrid strain were grown overnight in 3 ml cultures of YPD medium at 30°C. Upon confirming that the cultures were in log phase, two of the replicates were treated with 91 μl of DMSO, while the other two were treated with 91 μl of DMSO containing citrinin such that its final concentration in the culture was 600 ppm. Cultures were allowed to continue shaking at 30°C for two additional hours followed by harvesting of the pellets by centrifugation.

RNA was immediately isolated from the pellets using the MasterPure™ Yeast RNA Purification Kit (Epicenter). Two μg of RNA from each of the samples was subsequently used to create sequencing libraries using the low-throughput protocol in the TruSeq RNA Sample Preparation Kit v2 (Illumina). Individually barcoded libraries were pooled and sequenced (resulting in a total of ~156.6 million single-end 101 bp reads) on a single lane of an Illumina HiSeq 2000 instrument.

Reads were trimmed to 50 bp and mapped to a concatenated reference containing both parental genomes [29] using Bowtie version 0.12, allowing no mismatches [34]. Counts of reads mapping to a high-confidence curated ortholog set [35] were determined using htseq-count with the union option [36]. ASE was determined as the ratio of reads mapping to the *Sp* allele divided by those mapping to the *Sc* allele in the concatenated reference genome. Only genes with at least 20 reads mapping to each allele were retained. All reads are available in the NCBI SRA (accession PRJNA270666), and allele-specific read counts are given in Table S1.

To create the DVBPG6304/*Sp* hybrid, we started with diploid strains of each parent, and replaced one allele of the *URA3* gene with an antibiotic resistance gene (*hyg* in DVBPG6304 and *kan* in *Sp*). Transformants were selected on the corresponding antibiotic. The diploids were then sporulated to create haploids, and mixed to allow random mating. Hybrids were selected on plates containing hygromycin B and G418. RNA-seq was performed as described above, with ~27.3 million single-end 36 bp reads generated.

### Gene knockouts

The complete coding regions of five candidate genes were replaced with the *hphMX6* antibiotic resistance gene via PCR-mediated gene disruption [33] in *Sp.* Transformants were grown on hygromycin B (Sigma).

### Over-expression with CRISPR/Cas9

Molecular cloning was done with Gibson Assembly [37]. *E. coli* minipreps were performed with QIAprep Spin Miniprep Kits (Qiagen). Transformation and preparation of competent *E. coli* DH5α were prepared using Mix & Go *E. coli* Transformation Kit & Buffer Set and Zymo Broth (Zymo Research). Competent *S. cerevisiae* (strain BY4741) was prepared either by standard lithium acetate transformation protocols [33] or using Frozen-EZ Yeast Transformation II Kit (Zymo Research).

Two yeast constructs were obtained from ref. 38 via Addgene: p414-TEF1p-Cas9-CYC1t containing human codon optimized *Streptococcus pyogenes* Cas9 under control of the Tef1 *S. cerevisiae* promoter; and p426-SNR52p-gRNA.CAN1.Y-SUP4t, expressing a guide RNA under control of the SNR52 polIII promoter.

Catalytically inactive Cas9 (dCas9) was made by by introducing the D10A and H840A mutations [39]. We next PCR amplified the Gal4 activator and linker domains from the pDEST23 plasmid from the ProQuest Two-Hybrid System (Life Technologies), and fused it to the N-terminus of dCas9. We also obtained an alternative dCas9 activator utilizing four repeats of the minimal domain of the herpes simplex viral protein 16 (VP64) fused to the C-terminus of dCas9 [21]. We hypothesized that combining a Gal4 activator domain on the N-terminus with a C-terminal VP64 domain would produce a more potent activator than either activator fusion individually. To control for potential differences due to different codon optimizations, we cloned the VP64 activator domain from ref. 21 onto the dCas9 gene from ref. 38. We also cloned the dCas9-VP64 activator into the pRS416 vector under control of the Tef1 promoter so that all Cas9 fusions were expressed under the same plasmid and promoter backgrounds. To build the Gal4-dCas9-VP64 activator, we linearized the pRS414-Tef1-nNLS-Gal4-dCas9-Cyc1t plasmid and added VP64 to the C-terminal end via Gibson Assembly, followed by cloning into the pRS416 vector background.

We also made a gRNA plasmid from the p426-SNR52p-gRNA.CAN1.Y-SUP4t for cloning new guides by replacing the 20 base specificity sequence for the Can1.Y gRNA with a SacI restriction site. To activate the four candidate genes, we first designed two gRNAs for each gene. Because AG PAM sequences were reported to work as well as GG PAM sequences with dCas9 activators in human cell lines [40], we tested gRNAs with both type of PAM. Guides were selected to target between 50-250 bp upstream of the transcription start site (TSS) of each gene, because this window produced the best activation of other genes (data not shown) as well as in published data [21]. Guides were built from 60 base oligonucleotides, which contained 20 bases of overlap for Gibson cloning on either side of a 20 base unique specificity sequence. These single-stranded oligos were directly cloned into SacI digested pRS425-SacI-gRNA vector via Gibson assembly using an oligo to vector ratio of >100:1. This ratio was required because we found that Gibson Assembly less efficiently integrated single-stranded oligos than double-stranded fragments. PCR and Sanger sequencing were used to screen colonies. Confirmed clones were grown on LB carb media overnight and mini-prepped. These plasmids were then transformed into BY4741 containing the pRS416-dCas9-VP64 and grown on synthetic complete media plates lacking leucine and uracil for two days. From these plates, single colonies were selected and grown in liquid synthetic complete media lacking leucine and uracil. Activity of each gRNA was determined by qPCR. Interestingly, we found that none of the gRNAs designed with an AG PAM increased expression levels, suggesting that guides with an AG PAM may not be functional in *S. cerevisiae* (data not shown).

Once each gRNA was tested by qPCR, we sought to create a construct expressing a combination of guides, one to each of the four target genes. From each individual gRNA plasmid, we amplified the promoter, gRNA, and terminator using primers that produce unique overlaps to assemble the gRNAs in an ordered fashion. These PCR products were then assembled via Gibson Assembly in the absence of vector, followed by a second round of Gibson Assembly into pRS425. By doing a two-step Gibson Assembly, we drastically reduced the rate of misassembled products resulting from the high sequence similarity of the fragments. All constructs were verified by sequencing.

The four-gene over-expression strain in Fig. 4C was created by transforming the pRS416-Gal4-dCas9-VP64 plasmid and pRS425 with 4 gRNAs into BY4741. The control strain was identical to this, except using pRS425 with a nonfunctional gRNA.

### Strain Barcoding

Strains were barcoded by transforming a pRS416 vector (either empty pRS416 or pRS416-nGal4-dCas9-cVP64 for the dCas9-activator strains in Fig 4D) containing a 6-base barcode. These were constructed by integrating an oligo with a random 6mer in the center flanked by sequences matching our sequencing primers, and sequences matching the vector at the NsiI site for Gibson Assembly. Assembled vectors were sequence verified and then transformed into each strain of interest. All barcodes were at least two nucleotides separated from each other. For each strain, we transformed three unique barcodes. Each barcoded strain was then grown to saturation and mixed evenly with the other strains of that background and frozen as glycerol stocks.

### Competitive Growth Assays

Aliquots of the pooled barcoded strains were recovered in YPD media for 4 hours, and then diluted to OD 0.025 for the experiments. Yeast culturing and sample collection was performed using a cell-screening platform that integrates temperature-controlled absorbance plate readers, plate coolers, and a liquid handling robot. Briefly, 700 μl yeast cultures were grown in 48 well plates at 30° C with orbital shaking in either YPD media or YPD + 300ppm citrinin in Infinite plate readers (Tecan). To maintain cultures in log phase, 23 μl of the culture was removed when it reached an OD of 0.76, transferred to a well containing 700 μl of media, and then allowed to grow further. After seven such dilutions, 600 μl of the culture was collected at OD 0.76 and saved in a 4° C cooling station (Torrey Pines). This amounted to approximately 40 culture doublings from the beginning of the experiment. Fresh media transfers were triggered automatically by Pegasus Software and performed by a Freedom EVO workstation (Tecan).

After sample collection, yeast plasmids were purified using the Zymoprep Yeast Plasmid Miniprep II kit (Zymo Research). Purified plasmids were used as a template for PCR with barcoded sequencing primers that produce a double index to uniquely identify each sample. PCR products were confirmed by gel electrophoresis. After PCR, samples were combined and bead cleaned with Sera-Mag Speed Beads Carboxylate-Modified particles (Thermo Scientific). Sequencing was performed using Illumina MiSeq. Reads were counted only if they were a perfect match to the expected 6 bp barcode and flanking sequences.

### Promoter replacement strain construction

*In vivo* site-directed mutagenesis, known as *delitto perfetto* [22], was performed as described [40]. Briefly, the *Kluyveromyces lactis URA3* gene was amplified using primers containing ~70 bp of homology to each *Sc* promoter. This PCR product was transformed into *Sc,* and correct incorporation into each promoter was verified by PCR. Each *URA3* gene was then removed by transforming a PCR-amplified orthologous *Sp* promoter, containing enough flanking DNA sequence (identical between *Sc* and *Sp*) to allow specific targeting of the PCR product. Because the efficiency of *delitto perfetto* is maximized when transforming longer DNA molecules, as well as double-stranded DNA [33], transforming long PCR products (as opposed to shorter, single-stranded synthetic oligonucleotides) is a useful modification. Counter-selection of the resulting transformants on 5-FOA (Sigma) allowed isolation of successfully engineered strains that had replaced each *URA3* gene with the correct *Sp* promoter, which were then sequence-verified.

### Quantitative PCR

To measure the expression levels of our four candidate genes (Figs. 4C and 5B), we followed our previously published methods for qPCR [41]. We first grew each strain in YPD, and harvested them in log-phase (OD_600_ ~1) by centrifugation. We extracted total RNA with the MasterPure™ Yeast RNA Purification Kit (Epicentre), and quantified them with a NanoDrop2000 spectrophotometer. Total RNA samples were diluted to a concentration of 500 ng/μL and then reverse transcribed into cDNA with SuperScript III RT (Invitrogen), following manufacturer protocols. cDNA was diluted 1:100 prior to qPCR. qPCR was performed on an Eco Real-Time PCR machine (Illumina) following manufacturer’s protocols. To quantify changes in mRNA abundance, six control genes previously noted for their stability across conditions [42] were measured in each experiment: *ACT1, TDH3, ALG9, TAF10, TFC1,* and *UBC6.* All measurements were performed in triplicate. Data were analyzed using qBase Plus software (Biogazelle) [43].

## Acknowledgements

We would like to thank members of the Fraser Lab and the Stanford Genome Technology Center for discussions and advice. This work was supported by NIH grants P01HG000205 and 1R01GM097171-01A1. JDS is supported by a Genentech Graduate Fellowship. HBF is a Pew Scholar in the Biomedical Sciences.

---

**Figure S1. Reproducibility of biological replicates. A.** Reads per gene are shown for RNA-seq replicates of the *Sc/Sp* hybrid grown in YPD + DMSO. **B.** Reads per gene are shown for RNA-seq replicates of the *Sc/Sp* hybrid grown in YPD + 600 ppm citrinin (dissolved in DMSO).

**Figure S2. Agreement between RNA-seq and microarray data. A.** ASE ratios for the *Sc/Sp* hybrid in microarray data in YPD [16] compared to our RNA-seq data in YPD+DMSO. **B.** Fold-changes for *Sc’*s response to 300 ppm citrinin from oligonucleotide microarray data [15], compared to the *Sc* allele’s response (within the *Sc/Sp* hybrid) to 600 ppm citrinin in our RNA-seq data.

**Table S1. Read counts from RNA-seq in the Sc/Sp hybrid.**

**Table S2. All primers/oligos used in this work.**

